# Replacing bar graphs of continuous data with more informative graphics: Are we making progress?

**DOI:** 10.1101/2022.03.14.484206

**Authors:** Nico Riedel, Robert Schulz, Vartan Kazezian, Tracey Weissgerber

## Abstract

Recent work has raised awareness about the need to replace bar graphs of continuous data with informative graphs showing the data distribution. The impact of these efforts is not known. This observational meta-research study examined how often scientists in different fields use various graph types, and assessed whether visualization practices have changed between 2010 and 2020. We developed and validated an automated screening tool, designed to identify bar graphs of counts or proportions, bar graphs of continuous data, bar graphs with dot plots, dot plots, box plots, violin plots, histograms, pie charts, and flow charts. Papers from 23 fields (approximately 1,000 papers/field/year) were randomly selected from PubMed Central and screened (n=227,998). F1 scores for different graphs ranged between 0.83 and 0.95 in the internal validation set. While the tool also performed well in external validation sets, F1 scores were lower for uncommon graphs. Bar graphs are more often used incorrectly to display continuous data than they are used correctly to display counts or proportions. The proportion of papers that use bar graphs of continuous data varies markedly across fields (range in 2020: 4%-58%), with high rates in biochemistry and cell biology, complementary and alternative medicine, physiology, genetics, oncology and carcinogenesis, pharmacology, microbiology and immunology. Visualization practices have improved in some fields in recent years. Fewer than 25% of papers use flow charts, which provide information about attrition and the risk of bias. This study highlights the need for continued interventions to improve visualization and identifies fields that would benefit most.

## Introduction

The use of bar graphs to present continuous data remains common despite both meta-research studies [1, 2] and opinion pieces [3] highlighting the problems with this approach. Bar graphs are intended to display counts or proportions, where the bar is filled with data and a higher bar corresponds to a higher count or proportion. In contrast, bar graphs should not be used to present summary statistics (e.g. mean and standard error or standard deviation) for continuous data [1, 2]. Datasets with many different distributions may have the same summary statistics. The actual data may suggest different conclusions from the summary statistics alone. Continuous data should be presented using graphs that allow readers to examine the data distribution [1, 2]. Dot plots are best for small datasets, as summary statistics are only meaningful when there are enough data to summarize [1, 2]. Dot plots can be combined with box plots or violin plots for medium-sized datasets, while box plots, violin plots or histograms are recommended for larger datasets [2].

A 2015 paper [1] that was widely circulated by scientists in many fields raised awareness about the problems with bar graphs, prompting journals such as PLOS Biology [4], JBC [5] and eLife [6] to implement policy changes. New policies encouraged authors to replace bar graphs of continuous data with more informative graphics. More recently, other journals and publishers in various fields followed these early adopters by introducing their own policies (e.g. [7]). Researchers have also published free tools to help scientists create more informative graphics [3, 8-10]. Yet, a 2019 follow-up study [2] revealed that bar graphs of continuous data were the most common graph type among papers published in top peripheral vascular disease journals. Almost half of original research articles contained at least one bar graph of continuous data [2]. It is not know whether increased awareness and discussion about the problems with bar graphs has changed data visualization practices in the biomedical and biological sciences research communities.

Two other visualizations that have attracted attention within the scientific community are pie charts and flow charts. When displaying proportions, scientists can choose between pie charts, stacked bar charts and individual bar charts. In pie charts and stacked bar charts, slices of the pie or segments of the bar add up to 100% [11]. Pie charts require readers compare proportions by assessing angles and areas, whereas bar charts allow readers to compare lengths. Research suggests that bar charts of proportions are easier to interpret, as comparing lengths is easier than comparing angles and areas [12, 13]. Flow charts can be created for human, animal and in vitro studies, and report the number of included and excluded observations at each phase of the experiment [2]. These figures, which are recommended by some reporting guidelines (e.g. [14, 15]), provide essential information about attrition and the risk of bias.

This observational meta-research study sought to assess how often scientists in different fields use these various graph types, and to determine whether visualization practices have changed between 2010 and 2020. This will allow the scientific community to identify fields that might benefit from interventions to encourage authors to replace bar graphs of continuous data with more informative graphics. These data will also help us to determine whether data visualization practices have improved in recent years. The study was conducted using an automated screening tool, named Barzooka (RRID:SCR_018508), which we developed to identify the different graph types in manuscript PDFs. Screening was performed on 227,998 open access manuscripts deposited in PubMed Central.

## Methods

### Data availability statement

Open code for the trained algorithm for the Barzooka screening tool is deposited on Github (https://github.com/quest-bih/barzooka, RRID:SCR_018508, [16]). Open data and open code for the external validation analyses and the analyses examining the effects of field and time are available on the Open Science Framework [17].

### Automated screening tool development

Barzooka was trained to detect nine different types of graphs. These include: 1) bar graphs showing counts or proportions, 2) bar graphs that are inappropriately used to display continuous data, 3) bar graphs combined with dot plots, 4) dot plots, 5) box plots, 6) histograms, and 7) violin plots. The algorithm also detects pie charts, and flow charts that display the number of included and excluded observations at each stage of the study. Barzooka accepts a PDF of a publication as an input. Each page is converted to an image. The neural network then screens each page to determine whether the page includes graphs from each of the nine categories.Results from each page of the publication are aggregated to obtain publication-level data.

Barzooka was trained using the fastai v2.5.2 python library (RRID:SCR_022061) [18]. A pre-trained deep convolutional neural network (resnet101 pretrained on the ImageNet dataset) was trained using transfer learning. During transfer learning, the final layer of the original pre-trained network is replaced by a new layer, corresponding to the number of classes for the new classification task. The connections of the final layer are then retrained on data for different graph types, along with a few previous layers, which are trained at lower weights. This approach requires less training data because the information stored in the first layers of the pre-trained network, which is very generic and thus useful for any image classification task, can be reused.

The training dataset was derived from 14,000 open access publications downloaded from PubMed Central and eLife. Publications were split into pages. Each page was converted to jpeg images with a resolution of 560×560 pixels. The prediction was performed separately on each page (jpeg image). Training was based on between 1,000 and 5,000 manually labeled example pages per category. Single pages that included graphs from more than one category had multiple labels. This allows the network to predict several graph types per page. Some training pages did not contain graph types from any of the categories of interest. Pure text pages were labeled as ‘text’, while pages with an image or graph type not considered by the tool were labeled as ‘other’.

The network was trained using a batch size of 16 and ten training cycles. No further data augmentation (cropping, rotation, flipping) was used. This approach ensured that pages retained the standardized page format and layout, and that information on the pages was not altered. Ninety percent of pages in each category were used to train the network (n = 37,784), whereas 10% were used for validation (n = 3,812).

### bioRxiv validation set

The tool was validated on all preprints posted on bioRxiv between May 1 and May 15, 2019 (n = 1,107). Two independent reviewers (RS, VK) reviewed each preprint to determine whether the preprint contained graphs in each of the nine categories. Disagreements were resolved by consensus.

### Charité validation set

Additional validation for flow charts and pie charts, two categories that were uncommon in the bioRxiv validation set, was performed on articles published by authors affiliated with Charité Universitätsmedizin – Berlin or the Berlin Institute of Health at Charité, between 2015 and 2019. One thousand articles were randomly selected. Two independent reviewers (RS, VK) reviewed each preprint to determine whether the preprint contained graphs in each of the nine categories. Disagreements were resolved by consensus.

### Examining the effects of field and time

The dataset was derived from open access articles deposited in PubMed Central. This large dataset is easy to access. Original research articles were identified using the “article-type=research-article” tag in the PubMed Central XML file (https://www.ncbi.nlm.nih.gov/pmc/pmcdoc/tagging-guidelines/article/dobs.html#dob-arttype-resart). Other article types, such as reviews, editorials and comments, were excluded. We used the Fields of Research from the Australian and New Zealand Standard Research Classification [19] to determine article field, based on the journal in which the article was published. Under this system, 25,000 journals were classified into fields based on expert opinion. Each journal can be classified into a maximum of three fields. Journals that publish research in more than four categories are classified as multidisciplinary. We selected 23 fields, focusing on those that were particularly relevant for the biomedical and biological sciences. Some combinations of classifications were frequently assigned to the same journals. In these cases, we also sought to minimize the number of journals with multiple classifications by excluding one of these fields. Table S1 lists selected fields and codes.

Sampling was conducted over an 11-year period (2010 to 2020). We randomly selected 1,000 research articles per field per year. In smaller fields, where the sample did not contain 1,000 articles per year, all available articles were included (Table S2). The Fields of Research classification can assign up to three different fields to one journal. For articles in journals that are assigned multiple of the selected fields, one field is chosen at random during sampling. For sample generation, we used a dataset previously generated by Serghiou et al. [20] that combines PMC articles with additional metadata (e.g. article type). For newer PMC articles in our sample that were not yet included in this dataset, we retrieved the metadata separately. A total of 227,998 articles were screened using one graphics processing unit (GPU) node of the Berlin Institute of Health computing cluster.

PDFs were downloaded automatically using via the PMC FTP service using the direct PDF link or the compressed archives of the articles. In cases where no PDF could be found, we tried to obtain a PDF version via the unpaywall API or, for Elsevier articles, the Elsevier API. Some publishers include supplemental material at the end of article PDFs, whereas others do not. Supplemental files that were not included in the manuscript PDF were not screened.

### Data analysis and figure creation

The proportion of papers that used each graph type in each field and year were calculated. Precision for each graph type was calculated as the number of true positive cases divided by all cases detected by the algorithm. Recall was calculated as the number of true positive cases divided by all cases detected by the human raters. F1 scores were calculated as harmonic means of precision and recall for each class. Harmonic means are typically used when averaging rates or ratios. Graphs were prepared using the ggplot2 R package (RRID:SCR014601). As this was a descriptive study, no formal statistical analyses were performed. However, 95% confidence intervals are shown to estimate the precision of the observed effects.

## Results

### Internal validation

Table 1 shows performance data for the internal validation set. F1 scores ranged between 0.83 (histograms) and 0.95 (violin plots). Precision ranged between 0.84 (bar graphs of counts and proportions) and 0.97 (pie charts). Recall ranged between 0.80 (histograms) and 0.92 (pie charts).

**Table 1:**
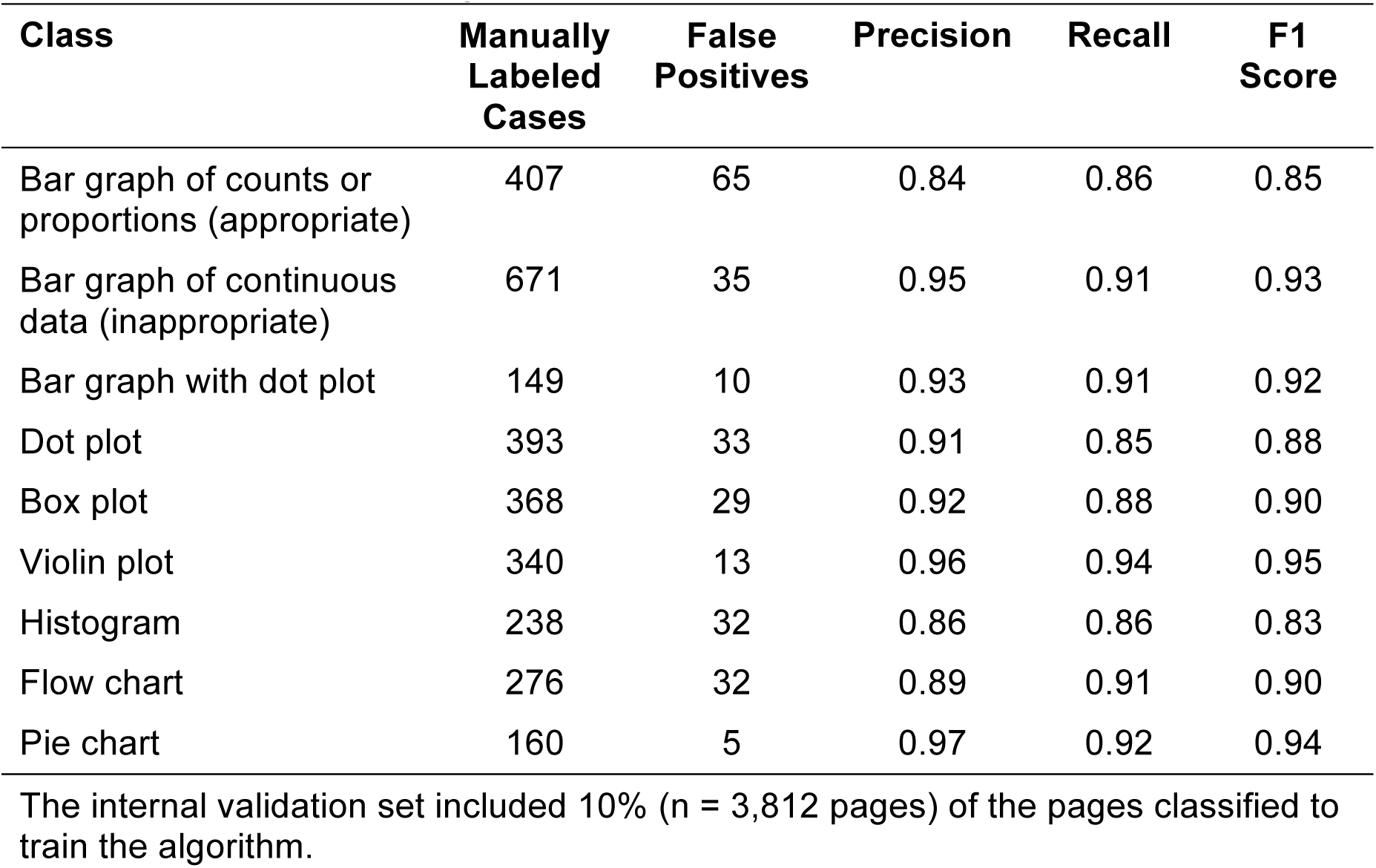
Barzooka screening tool performance in an internal validation set

### bioRxiv validation set

The tool was further validated on a set of 1,107 preprints published on bioRxiv between May 1 and May 15, 2019. While the tool was still effective, F1 scores, precision and recall were lower than in the internal validation set (Table 2). Performance was lowest in categories that were uncommon in the dataset. In these categories, the number of false positives is high compared to the number of cases, which lowers F1 scores.

**Table 2:** Barzooka screening tool performance an external validation set of bioRxiv preprints

| **Class** | **Manually Labeled Cases** | **False Positives** | **Precision** | **Recall** | **F1 Score** |
| --- | --- | --- | --- | --- | --- |
| Bar graph of counts or proportions (appropriate) | 345 | 60 | 0.82 | 0.81 | 0.82 |
| Bar graph of continuous data (inappropriate) | 405 | 25 | 0.94 | 0.92 | 0.93 |
| Bar graph with dot plot | 74 | 37 | 0.63 | 0.86 | 0.73 |
| Dot plot | 257 | 51 | 0.80 | 0.80 | 0.80 |
| Box plot | 255 | 36 | 0.87 | 0.91 | 0.89 |
| Violin plot | 57 | 27 | 0.65 | 0.89 | 0.76 |
| Histogram | 198 | 66 | 0.72 | 0.85 | 0.78 |
| Flow chart | 20 | 26 | 0.40 | 0.85 | 0.54 |
| Pie chart | 71 | 12 | 0.83 | 0.85 | 0.84 |

The bioRxiv external validation set included 1,107 bioRxiv preprints published in May 2019.

### Charité validation set

Additional validation for flow charts and pie charts, two categories that were uncommon in the bioRxiv validation set, was performed on a set of 1,000 articles published by authors affiliated with Charité Universitätsmedizin – Berlin or the Berlin Institute of Health at Charité. Flow charts and pie charts were more common in this dataset, resulting in higher F1 scores of 0.88 and 0.86 (Table 3).

**Table 3:** Barzooka screening tool performance an external validation set of publications from authors affiliated with Charité Universitätsmedizin - Berlin

| **Class** | **Manually**<br>**Labeled**<br>**Cases** | **False**<br>**Positives** | **Precision** | **Recall** | **F1**<br>**Score** |
| --- | --- | --- | --- | --- | --- |
| Flow chart | 123 | 20 | 0.84 | 0.87 | 0.86 |
| Pie chart | 38 | 4 | 0.89 | 0.87 | 0.88 |

The Charité external validation set included 1,000 papers published between year and year.

### Sample generation

Figure 1 shows the number of excluded observations and reasons for exclusion. 3,299 (1.4%) articles in the sample were excluded because full text PDFs could not be downloaded automatically. A total of 227,998 articles open access articles published between 2010 and 2020 were screened.

**Figure 1:**
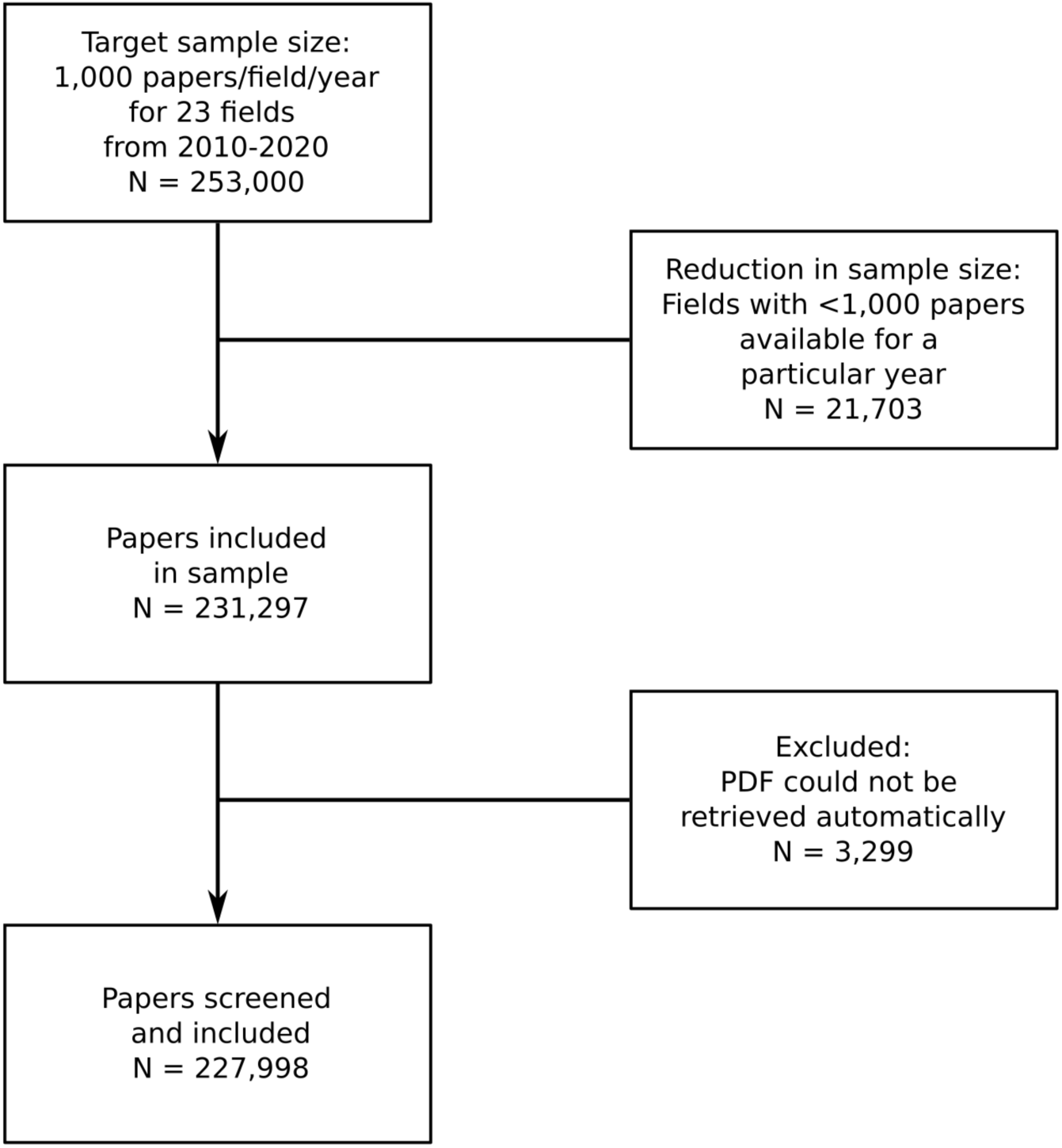
Flow chart The flow chart illustrates the sampling strategy and shows the number of papers excluded and reasons for exclusion at each phase of the experiment.

### Bar graphs

The proportion of papers that use bar graphs to display continuous data was highly variable across different fields, ranging from approximately 5% to 60% of papers (Figure 2). Bar graphs of continuous data are common in fields such as biochemistry and cell biology, physiology, genetics, complementary and alternative medicine and oncology and carcinogenesis. In contrast, bar graphs of continuous data were uncommon in fields like nursing, dentistry and pediatrics and reproductive medicine.

**Figure 2:**
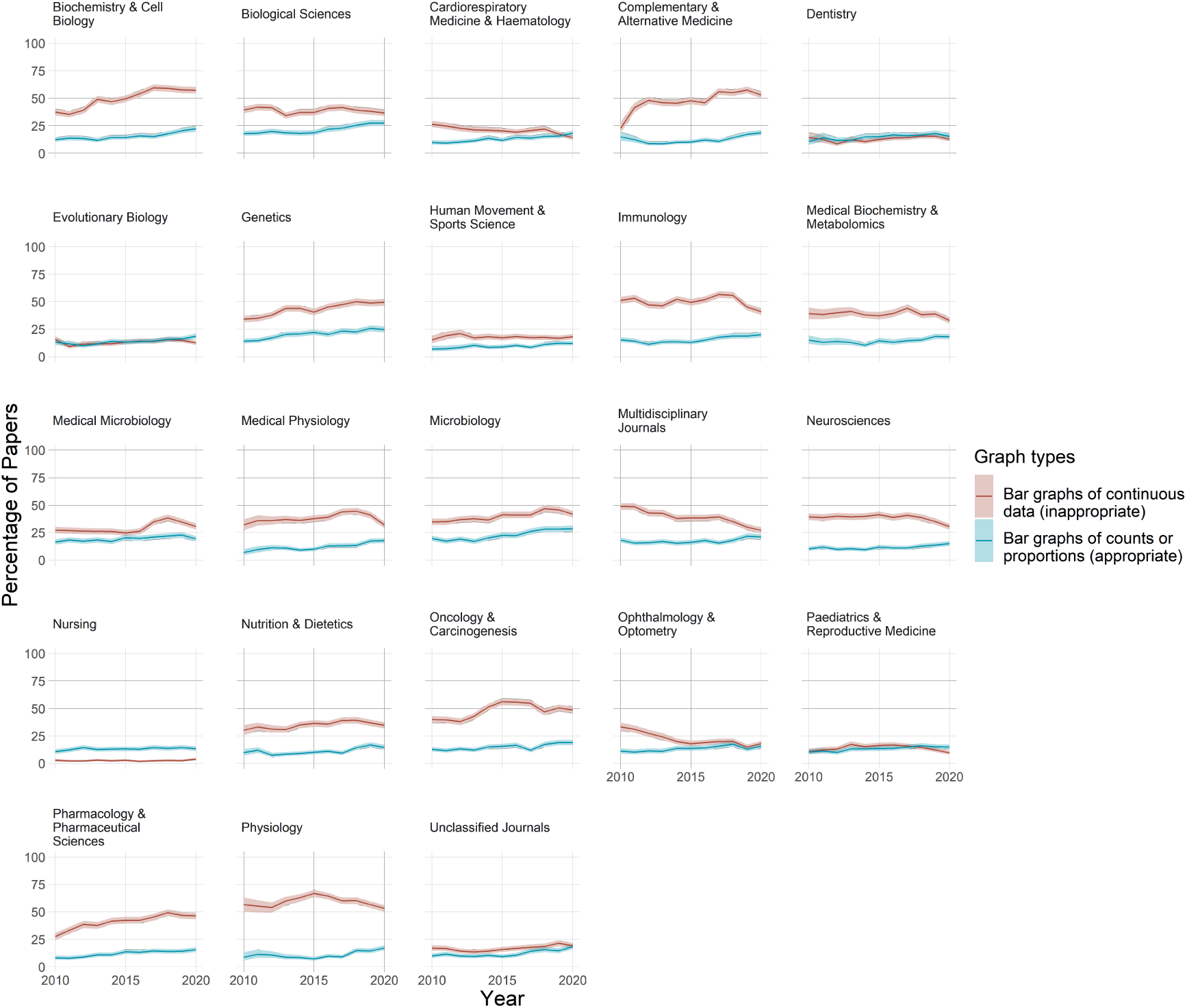
Bar graphs are commonly used incorrectly Small multiples show the percentage of papers that use bar graphs to display continuous data (red line), vs. counts and proportions (blue line), for 23 fields over an 11-year period. Shaded regions above and below each bar display 95% confidence intervals. The percentage of papers that use bar graphs for continuous data ranges between approximately 5% and 60% of papers, depending on the field. With the exception of a few fields where bar graphs are uncommon (e.g. nursing, dentistry), the improper use of bar graphs to display continuous data is more common than the correct use of bar graphs to display counts and proportions.

The inappropriate use of bar graphs to display continuous data is more common than the appropriate use of bar graphs to display counts and proportions (Figure 2). The proportion of papers that use bar graphs to display counts or proportions is consistently below 25% in almost all fields. In contrast, the proportion of papers that used bar graphs to display continuous data was between 30 and 60% for 15 out of 23 fields.

The use of bar graphs of continuous data appeared to be increasing over time in fields like biochemistry and cell biology, genetics, complementary and alternative medicine and pharmacology and toxicology. In other fields, bar graph use was relatively stable between 2010 and 2020 (e.g. biological sciences, evolutionary biology, human movement and sport sciences). In recent years, the proportion of papers that use of bar graphs of continuous data appeared to be decreasing in fields such as immunology, physiology, medical physiology, neuroscience, and multidisciplinary journals.

Among all papers that were screened, the proportion of papers that included bar graphs of continuous data may have decreased slightly in recent years, while the proportion of papers that combine bar graph with dot plots may have increased slightly (Figure 3). The proportion of papers that include more informative graphs, specifically dot plots, box plots, violin plots, or histograms, appears to have increased by more than 10%. Note that these results apply only to the sample screened, which does not reflect the proportion of papers in each field available in PubMed Central.

**Figure 3:**
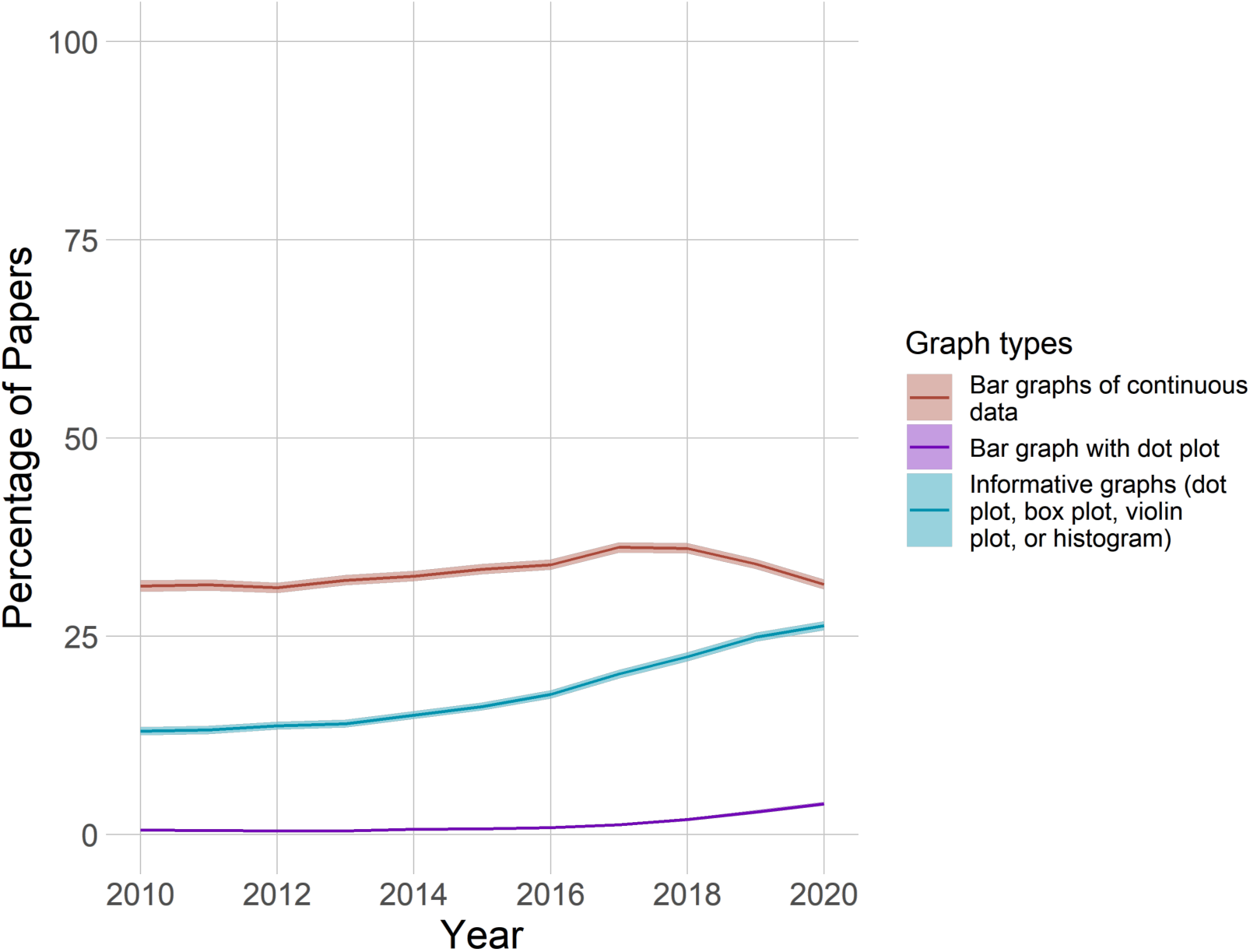
Time trends for visualizing continuous data in the complete sample The graph shows the percentage of papers with bar graphs of continuous data (red), bar graphs with dot plots (purple) and more informative graphs (dot plots, box plots, violin plots, or histograms; blue), from 2010 to 2020, in the complete sample (n = 227,998 papers). Shading shows 95% confidence intervals. The confidence intervals for bar graphs with dot plots are too small to be seen in the figure. These findings from the current sample should not be extended to all of PubMed Central, as the proportion of articles in different fields differs between the study sample and PubMed Central.

### Bar graphs with dot plots

Bar graphs combined with dot plots are rare, occurring in less than 5% of papers in most fields (Figure 4). Despite these low baseline rates, use of these plots seems to have risen over the past few years in fields such as biochemistry and cell biology, physiology, medical physiology, immunology and neuroscience.

**Figure 4:**
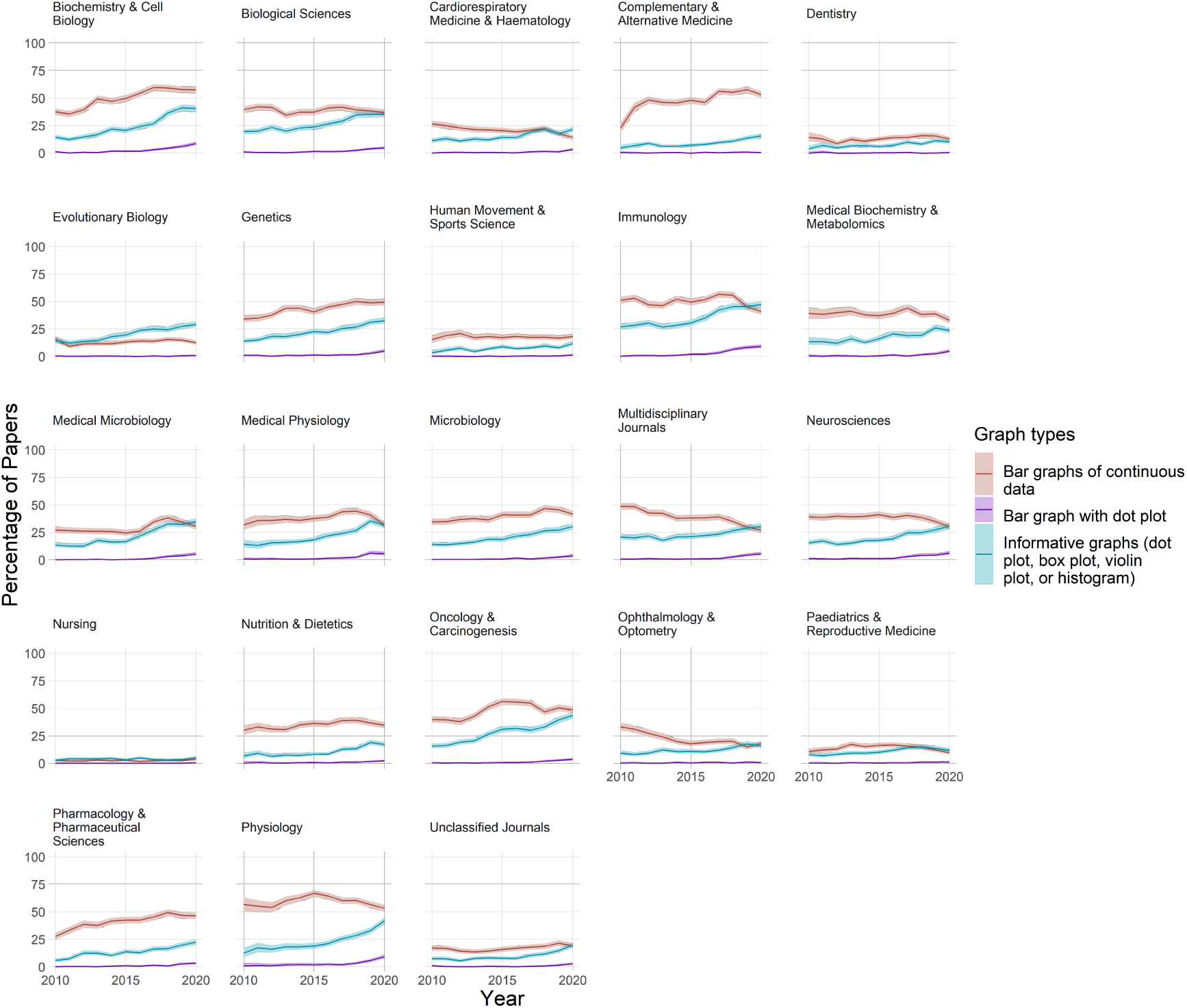
The use of bar graphs, bar graphs with dot plots, and more informative graphs to display continuous data Small multiples show data for 23 fields over an 11-year period. Lines show the percentage of papers that use bar graphs (red line), bar graphs with overlaid dot plots (purple line), or a more informative graph type (blue line) to display continuous data. Shaded regions show 95% confidence intervals. The informative graphs category includes dot plots, box plots, violin plots, and histograms. Bar graphs are the least informative graph type. Bar graphs with dots are suboptimal.

### Dot plots, box plots, violin plots and histograms

More informative alternatives to bar graphs of continuous data, including dot plots, bot plots, violin plots, and histograms, are less common than bar graphs in most fields (Figure 4). Use of these alternative graphics appears to be rising over time in most fields where bar graphs are commonly used. These changes appear to be driven primarily by increased use of dot plots and box plots (Figure 5). Violin plots are uncommon in most fields.

**Figure 5:**
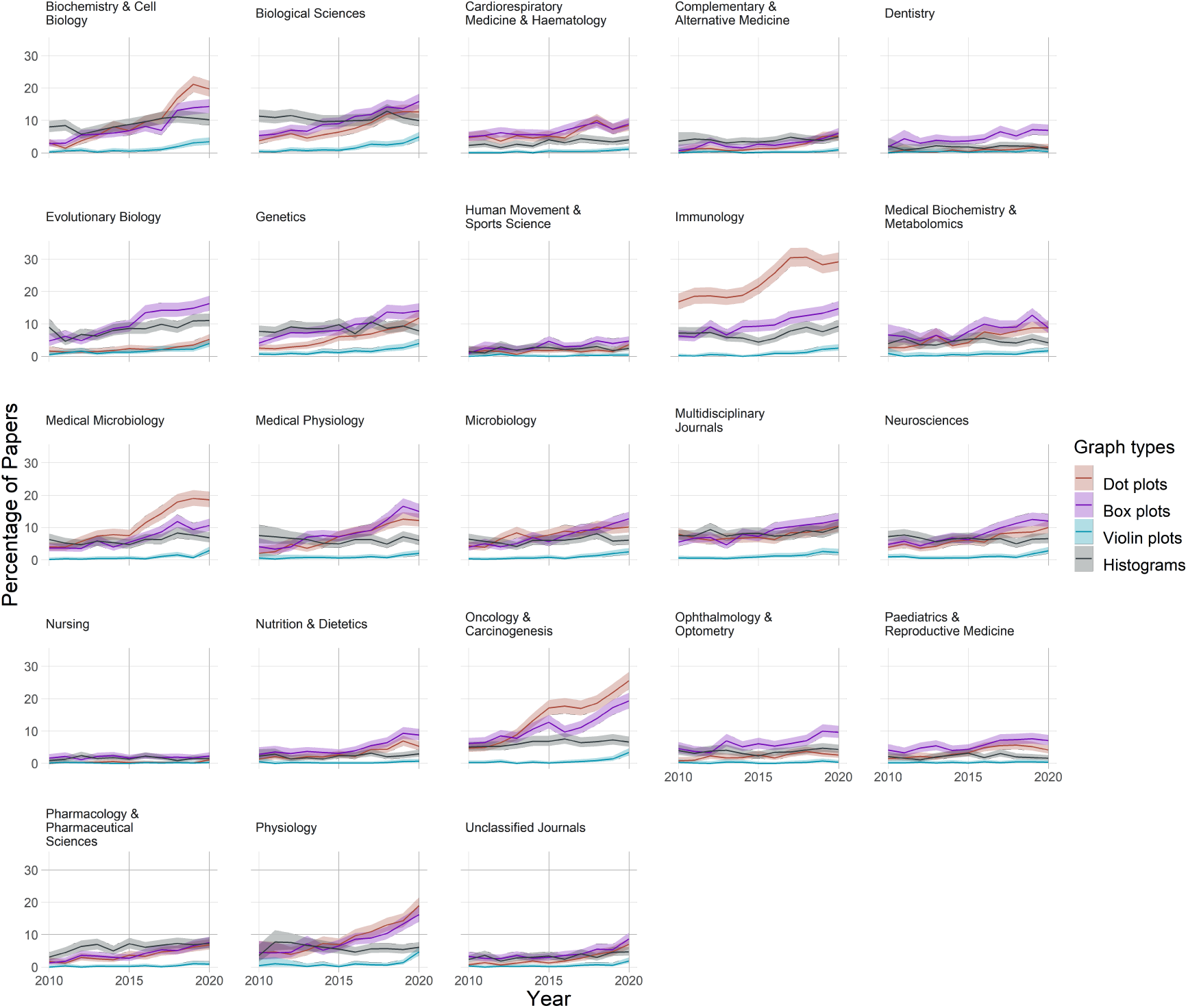
The use of bar graphs, bar graphs with dot plots, and more informative graphs to display continuous data Small multiples show data for 23 fields over an 11-year period. Lines show the percentage of papers that use dot plots (red line), box plots (purple line), violin plots (blue line) or histograms (gray line). Note that the y-axis shows values up to 30% to allow readers to see percentages for these less common graph types.

### Identifying fields that may benefit from interventions

We classified fields into three priority groups for interventions to encourage authors to replace bar graphs of continuous data with more informative graphics (Figure 6). Fields in which more than 40% of papers contain bar graphs of continuous data have the highest priority for interventions. These include microbiology, immunology, pharmacology and pharmaceutical sciences, genetics, complementary and alternative medicine, oncology and carcinogenesis, physiology and biochemistry and cell biology. In two of these fields, complementary and alternative medicine and pharmacology and pharmaceutical sciences, more informative graphs are not commonly used. Interventions in these fields may need to provide more information about how to determine what type of more informative graphs is best for one’s dataset, and how to make more informative graphs. Seven fields have a medium priority for interventions, with 20-40% of papers containing bar graphs of continuous data. These include multidisciplinary journals, neurosciences, medical microbiology, medical physiology, medical biochemistry and metabolomics, nutrition and dietetics, and biological sciences. The lowest priority for intervention is fields in which fewer than 20% of papers include bar graphs of continuous data. This category includes nursing, pediatrics and reproductive medicine, dentistry, cardiorespiratory medicine and hematology, evolutionary biology, human movement and sports science, ophthalmology and optometry and unclassified journals.

**Figure 6:**
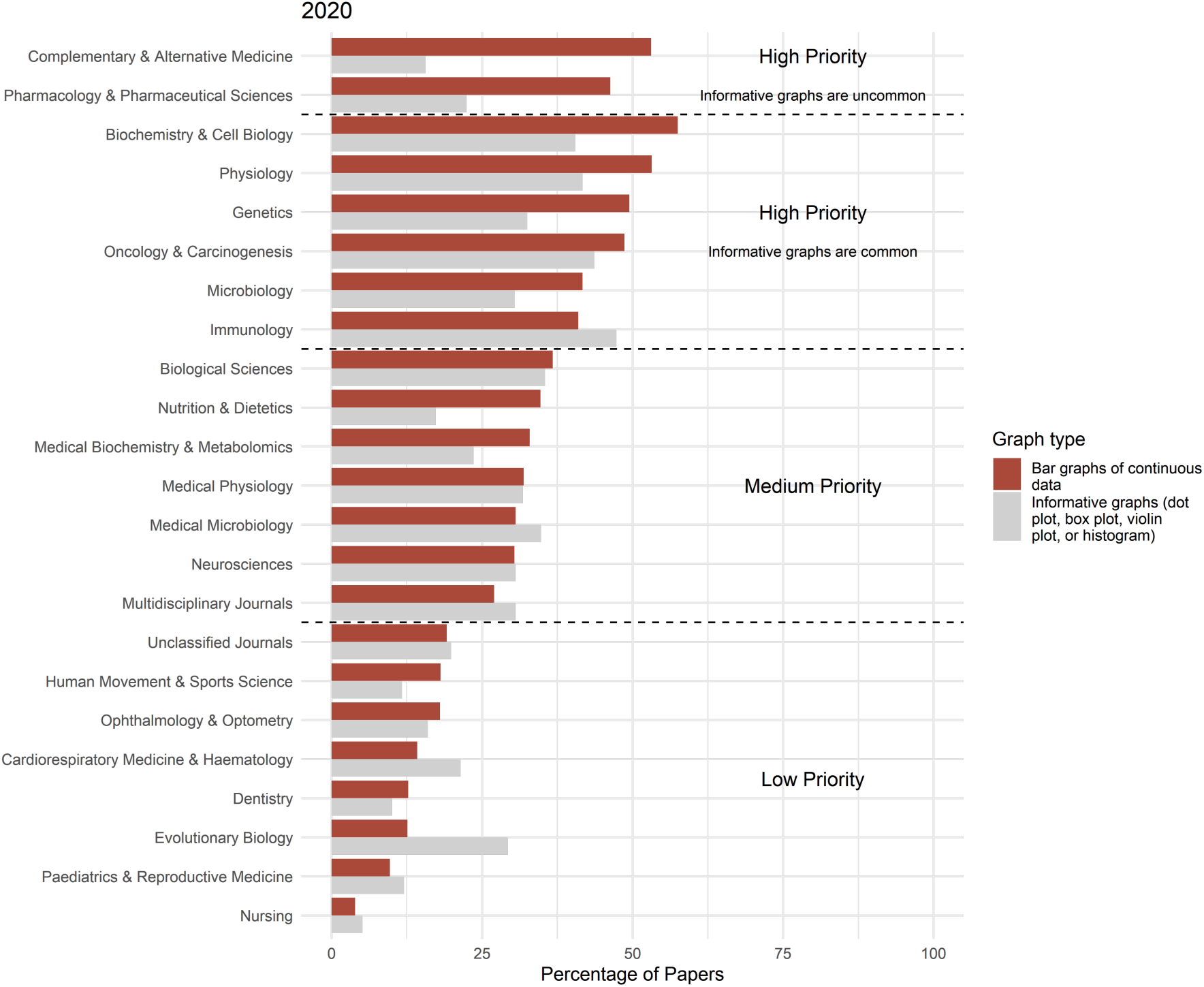
Identifying fields that would benefit from interventions Fields are divided into three priority groups, depending on the percentage of papers that use bar graphs to present continuous data (red bars; High priority: >40%; Medium priority: 30-40%; Low priority: <20%). The high priority group includes two subgroups; one subgroup in which more informative graph types are less common, and a second subgroup in which more informative graph types are more commonly used. Training in how to create more informative graphs, and how to choose between dot plots, box plots, violin plots and histograms may be especially critical for fields where more informative graphs are less common.

We next sought to determine whether authors who were using bar graphs of continuous data were also using more informative graphics. Small multiples showing the percentage of papers using only bar graphs, only more informative graphs, or a combination of bar graphs and more informative graphs, are shown in Figure 7. Bar graphs with dot plots were not included in this analysis, as they were rare in most fields. In some fields, the percentage of papers using only bar graphs appears to be falling in recent years, while the percentage of papers that use more informative graphs appears to be rising. Examples include physiology, immunology and neurosciences. While not conclusive, these data are consistent with the idea that some authors in these fields may be switching from bar graphs of continuous data to more informative graphs. These changes were not apparent in other fields, such as genetics and microbiology.

**Figure 7:**
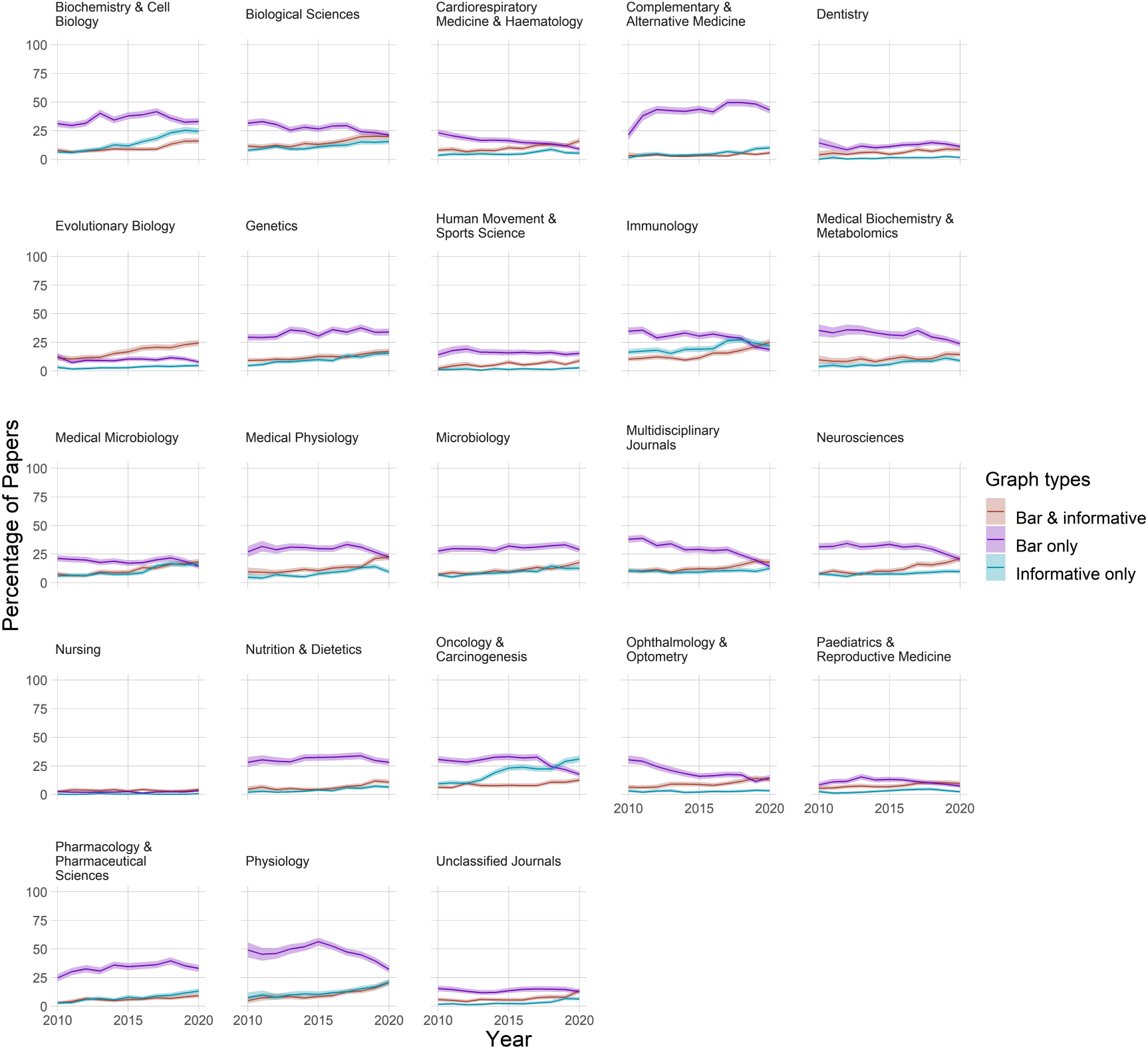
Are authors who use bar graphs of continuous data also using more informative graphs? Small multiples display the proportion of papers that include both bar graphs of continuous data and more informative graphs (dot plots, box plots, violin plots or histograms) (red line), only bar graphs of continuous data (purple line), or only more informative graphs (blue line) between 2010 and 2020. Shading represents the 95% confidence interval.

### Flow charts

Flow charts are uncommon, although use seems to be increasing over time in fields such as medical biochemistry and metabolomics, cardiorespiratory medicine and hematology, human movement and sport sciences, pediatrics and reproductive sciences and medical physiology (Figure 8). Even in these fields, the proportion of papers with flow charts in recent years remains below 25%. Fewer than 5% of papers have flow charts in fields such as the biological sciences, biochemistry and cell biology, evolutionary biology, genetics, microbiology and immunology.

**Figure 8:**
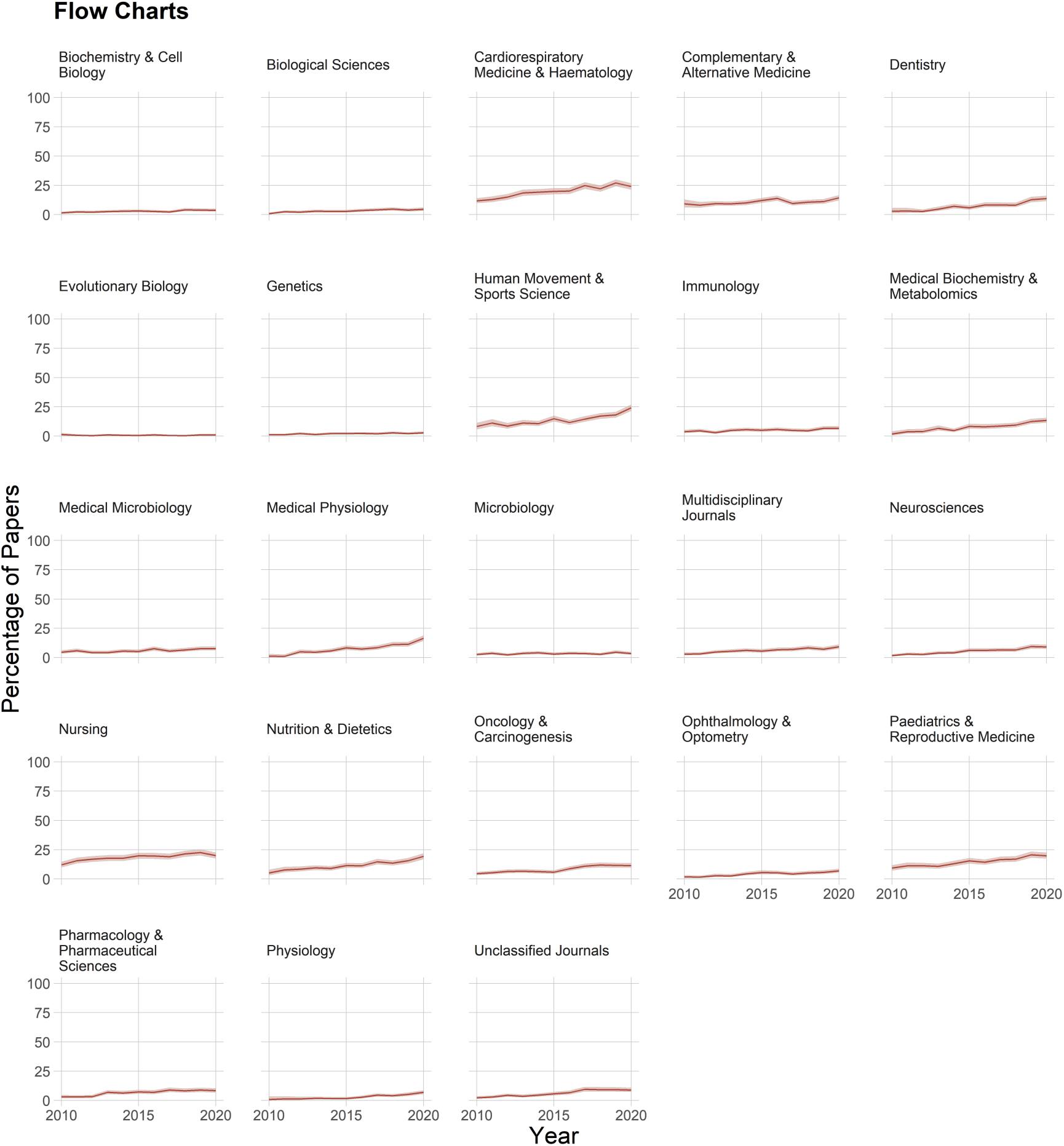
The use of flow charts Fewer than 25% of papers included flow charts. Shading shows 95% confidence intervals.

### Pie charts

Pie charts are also rarely used, appearing in fewer than 5% of papers in most fields (Figure 9).

**Figure 9:**
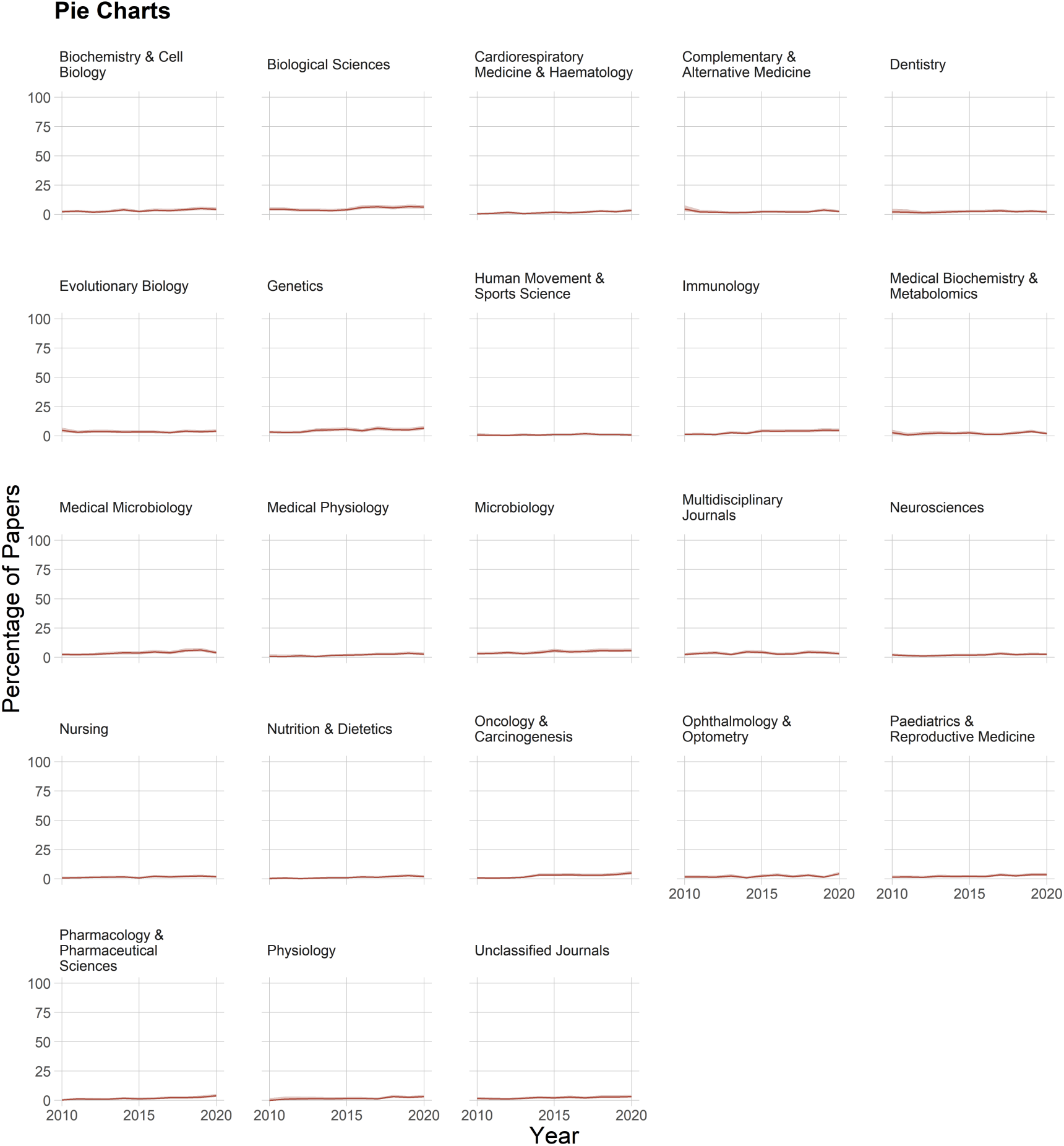
The use of pie charts Pie charts were rare in all fields. Shading shows 95% confidence intervals.

## Discussion

This research offers several valuable contributions. First, we introduce a new automated tool that can screen large numbers of publications for common graph types. Second, the data show that in many fields, bar graphs are more likely to be used incorrectly to display continuous data than they are to be used correctly to display counts and proportions. Third, results show that the proportion of papers that use bar graphs of continuous data varies markedly across fields. Interventions and educational initiatives encouraging authors to replace bar graphs of continuous data with more informative graphics should prioritize fields where bar graphs are commonly used. Fourth, trends in some fields suggest that the use of bar graphs of continuous data may have decreased in recent years, while the use of more informative alternative graph types may have increased. Further monitoring over the next few years is needed to determine whether these trends are early indicators of widespread change. If these trends are confirmed, researchers should determine whether these changes are be due to increasing awareness of the problems with bar graphs, journal policy changes, or other factors. Finally, the data highlight the need to encourage scientists to use flow charts to provide information about attrition and to help readers assess the risk of bias.

### Appropriate use of bar graphs

A previous study of papers published in top peripheral vascular disease journals in 2018 also found that bar graphs of continuous data were more common than bar graphs of counts or proportions [2]. The present study extends these findings to many other fields, while showing that this pattern was consistently observed from 2010 to 2020. Recent research indicates that some readers conflate bar graphs that show counts or proportions with bar graphs that show means of continuous data [21], even though these two data types have very different properties. When examining a bar graph of continuous data, one in five people incorrectly interpret the bar end as the maximum of the underlying data points, rather than the center of the data points [21]. This misinterpretation was found across general education levels [21]. An earlier study found that when comparing equidistant data points above and below the bar tip, viewers rated points within the bar as being more likely than points above the bar [22]. These two studies suggest that bar graphs are not as simple as they seem, and may be misinterpreted.

A recently developed automated screening tool identifies bar graphs in which the y-axis does not start at zero [23]. Our data highlight that it would be best to apply this tool only to bar graphs of counts or proportions. When bar graphs are used to display continuous data, authors should instead be encouraged to replace the graph with a dot plot, box plot or violin plot. Furthermore, starting the y-axis at zero is much more important for counts or proportions than for continuous data (Figure 10). Cutting the y-axis may be misleading for bar graphs of counts or proportions, as it alters viewers’ perception of the count or proportion. When graphing continuous data, however, there are clear cases where one should not start the y-axis at 0. When the dataset includes negative values, starting the y-axis at 0 would be misleading. When the “Zone of Irrelevance” (area at the bottom of the graph where there are no data points) [10] is very large, starting the y-axis at 0 may interfere with readers’ ability to examine the data distribution and overlap between groups. Starting the y-axis at 0 is most effective when all values are positive and the “Zone of Irrelevance” does not occupy a large portion of the graph. When graphing continuous data, scientists should choose a scale that makes sense for their dataset. Researchers can visually emphasize axis cuts by using wavy axes or cut lines that extend across the entire x-axis.

**Figure 10:**
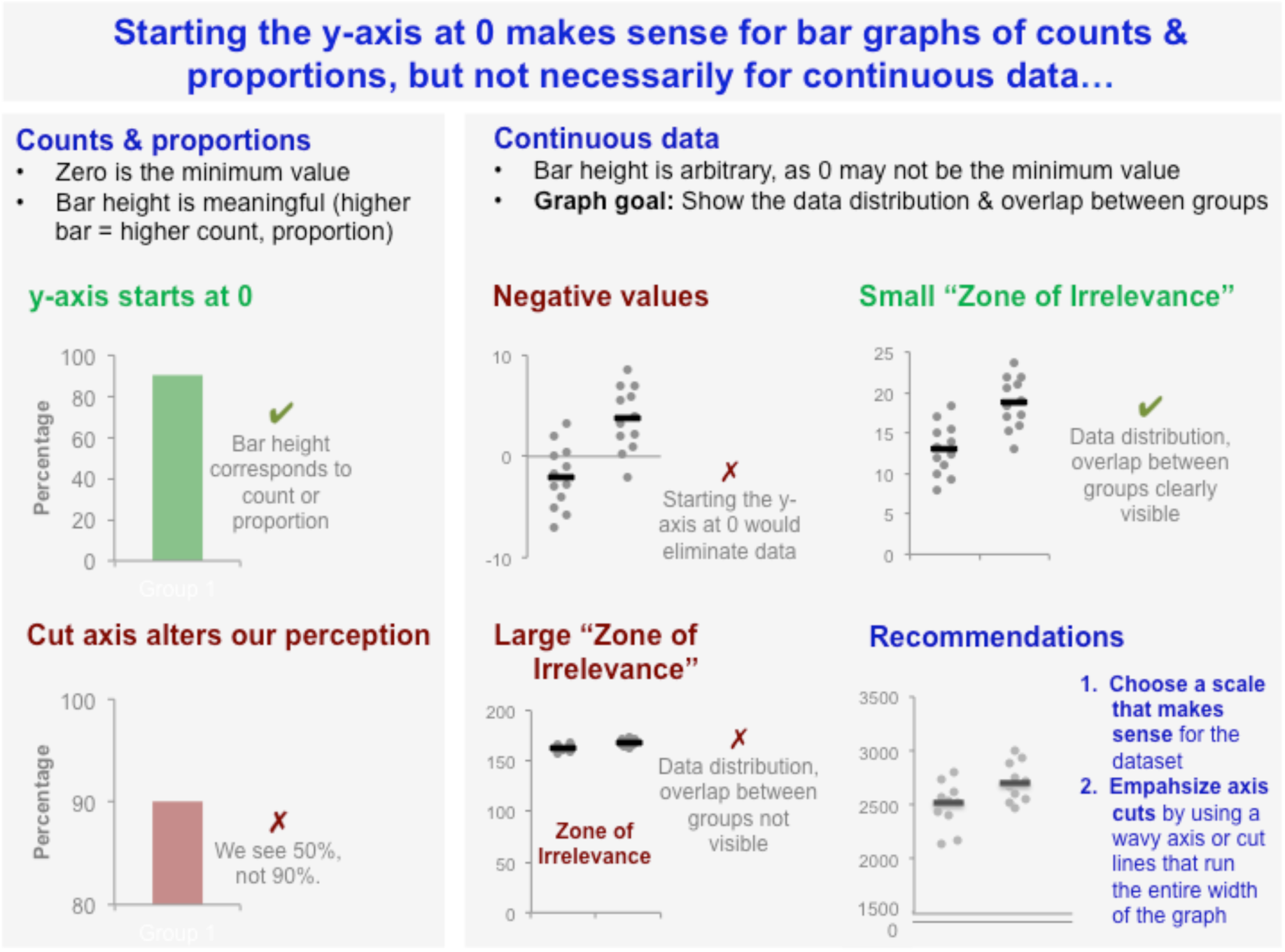
Starting the y-axis at 0 makes sense for bar graphs of counts and proportions, but not for continuous data This figure examines how the differing characteristics of counts and proportions vs. continuous data, effect decisions about whether to start the y-axis at 0. Cutting the axis is potentially problematic for counts and proportions, as this alters viewers’ perception of the count or proportion. In contrast, selecting a y-axis scale that does not start a zero may be essential for continuous data if the dataset includes negative values, or much of the graph is taken up by the “Zone of Irrelevance”. The “Zone of Irrelevance” refers to a region at the bottom of the graph that does not contain data points.

While bar graphs with dot plots are rare, our data suggest that the proportion of papers using these graphs has risen slightly in the past few years. These graphs should be replaced with dot plots. The bars are chart junk – they create the illusion of certainty without adding information. The summary statistics shown by the bars appear equally certain whether the bar contains 2 data points or 2,000 data points, whereas readers should have much more confidence in the summary statistics derived from the larger dataset. Summary statistics could be displayed more efficiently using a mean or median line with error bars showing a standard deviation or 95% confidence interval, overlaid on a dot plot. Shaded bars may obscure data points within the bar. In contrast, dot plots efficiently illustrate the sample size, variability in the data and amount of overlap between groups without introducing chart junk or distortions.

### Identifying fields that would benefit from interventions

This research identifies many fields where bar graphs are commonly used incorrectly. These fields would benefit from interventions that encourage authors to replace bar graphs of continuous data with more informative graphics. Further research is needed to determine what types of interventions are most effective, however options may include training editors and peer reviewers, offering data visualization workshops and webinars in collaboration with scientific societies in these fields, or integrating manual or automated screening into the manuscript submission process. Studies examining the effectiveness of journal policies asking authors to replace bar graphs of continuous data with more informative graphics have not yet been conducted. However, research suggests that other types of journal policy changes have little impact on reporting, especially if policy changes are not accompanied by an implementation or enforcement plan [24-26].

### Flow charts

The absence of flow charts in most papers is a missed opportunity. Information about the number of excluded observations, and reasons for exclusion, is essential for assessing the risk of bias [27, 28]. Biased exclusion of observations increases the likelihood of spurious findings, especially in small sample size studies [28]. Among preclinical studies using animal models to investigate cancer and stroke, 7-8% of papers excluded observations without explanation [28]. Approximately two thirds of papers did not contain enough information to determine whether observations were excluded without explanation [28]. A small exploratory analysis suggested that clinical trials that included flow charts were more likely to fully report attrition than studies that provided this information in the text [29].

Observations are sometimes excluded for reasons that do not introduce bias. Awareness of all excluded observations can help scientists to plan future experiments, allocate resources, or improve their protocols. Knowing that only 50% of donor samples will be of sufficient quality for in vitro experiments, for example, allows scientists to more accurately estimate the number of required donors. This information might also prompt investigators to refine their sample collection or preparation protocols to increase the proportion of high quality samples.

Reporting guidelines for clinical trials [27] and systematic reviews [14] include flow chart templates. These charts can be adapted to other study designs. The Experimental Design Assistant (RRID:SCR_017019) generates flow charts for preclinical animal experiments [30]. While flow charts are not commonly used for in vitro research, templates from other types of studies may help investigators to design flow charts for to report attrition for individual experiments. Flow charts for studies using material from human or animal donors should start with the selection of donors and collection of donor material.

### Automated screening

Barzooka and other automated screening tools are valuable for both meta-research and interventions. Automated screening tools allow for a rapid assessment of articles. Barzooka needs only few seconds to screen a 20-page article when running on a graphics processing unit. While this study focused on meta-research, Barzooka is also part of the ScreenIT pipeline, which uses a set of automated tools to screen COVID-19 preprints posted on bioRxiv and medRxiv [31]. Public reports on new prepreints are posted on the website annotation tool Hypothes.is (RRID:SCR_000430) and a link is tweeted out via @SciScoreReports. Reports are also listed on Sciety, which aggregates preprint reviews. More than 18,000 COVID-19 preprints have been screened. While we hope that the reports will help authors to improve reporting before their preprint is published, intervention studies are needed to determine whether this approach is effective. Research is also needed to ensure that reports are useful to authors, and to determine how to best deliver reports to authors. Barzooka is also used to assess data visualization practices among papers published by authors at the Charité Universitätsmedizin – Berlin or the Berlin Institute of Health at Charité, and results are displayed on an institutional dashboard (https://quest-dashboard.charite.de). Some journals and publishers have used commercial automated screening tools, such as SciScore (RRID:SCR_016251) and Statcheck (RRID:SCR_015487), to screen submitted manuscripts [32, 33]. The process of integrating tools into manuscript submission systems is time consuming and very expensive. Solutions to efficiently and inexpensively integrate small open access tools into publishing systems are needed.

### Limitations

This study includes both technical limitations of the tool, and study design limitations. Barzooka may have difficulty distinguishing between bar graphs of continuous data, and the subset of bar graphs of counts or proportions that include error bars showing 95% confidence intervals. Visually, these graphs are very similar. The effects of field and time were examined in open access publications deposited in PubMed Central. These papers may systematically differ from paywalled papers published in subscription journals. Papers for 2020 were selected in January 2022, as funders such as the US National Institutes of Health have policies requiring that open access versions of all NIH-funded papers must be available in PubMed Central one year after publication [34]. While open access journals such as PLOS Biology [4] and eLife [6] have implemented policies encouraging authors to replace bar graphs with more informative graphics, some subscription journals have also implemented such policies [5, 7]. Papers from these journals are likely underrepresented in our sample, although previous research on other topics suggests that journal policy changes are not very effective [24-26]. Field codes were based on the journal in which the paper was published, rather than the content of the article itself. Reliable and effective strategies for determining field based on article content are needed. The tool may have systematically undercounted flow charts. Some authors place flow charts in the supplement and supplemental materials that were not included in the PDF of the paper were not screened.

## Conclusions

This study highlights the need for continued widespread interventions to improve data visualization practices across many scientific fields. Editors, reviewers and authors should carefully scrutinize all bar graphs, and ask authors to replace bar graphs of continuous data with more informative graphs. Dot plots should be used for small datasets, whereas dot plots may be combined with box plots or violin plots for medium-sized datasets. Box plots, violin plots, or histograms may be used for larger datasets. Bar graphs combined with dot plots should be replaced with more informative graphs. These actions are especially important for fields where bar graphs are often misused. Editors, reviewers and authors should strongly encourage authors to use flow charts for human, animal and in vitro studies. These figures convey essential information about attrition, lost observations, and the risk of bias.

Future studies should investigate the effectiveness of interventions to improve data visualization practices on a large scale, targeting fields where problems are most common. This may include studies evaluating the effectiveness of automated screening, journal policy changes or widespread educational initiatives. Creation of new automated screening tools may help the scientific community to explore the prevalence of other data visualization problems and conduct future interventions.

