## Supplemental_Tables for "Replacing bar graphs of continuous data with more informative graphics: Are we making progress?"

**Table S1:** Fields of Research classifications included in the study

| **Field** | **Code** |
| --- | --- |
| Biological sciences | 600 |
| Biochemistry & cell biology | 601 |
| Evolutionary biology | 603 |
| Genetics | 604 |
| Microbiology | 605 |
| Physiology | 606 |
| Medical biochemistry & metabolomics | 1101 |
| Cardiorespiratory medicine & Hematology | 1102 |
| Complementary & alternative medicine | 1104 |
| Dentistry | 1105 |
| Human movement & sport science | 1106 |
| Immunology | 1107 |
| Medical microbiology | 1108 |
| Neurosciences | 1109 |
| Nursing | 1110 |
| Nutrition & dietetics | 1111 |
| Oncology & carcinogenesis | 1112 |
| Ophthalmology & optometry | 1113 |
| Pediatrics & reproductive medicine | 1114 |
| Pharmacology & pharmaceutical sciences | 1115 |
| Medical physiology | 1116 |
| Multidisciplinary journals (≥4 categories) | N/A |
| Uncategorized journals | N/A |

Abbreviations: N/A, not applicable.

**Table S2:** Papers screened by field and year

| **Field** | **2010** | **2011** | **2012** | **2013** | **2014** | **2015** | **2016** | **2017** | **2018** | **2019** | **2020** |  | **Scale** |
| --- | --- | --- | --- | --- | --- | --- | --- | --- | --- | --- | --- | --- | --- |
| Biochemistry & Cell Biology | 993 | 987 | 989 | 966 | 996 | 996 | 994 | 998 | 1000 | 994 | 1000 |  | 200s |
| Biological Sciences | 999 | 994 | 999 | 992 | 995 | 999 | 996 | 998 | 996 | 992 | 996 |  | 300s |
| Cardiorespiratory Med. & Haematology | 891 | 789 | 887 | 875 | 968 | 924 | 997 | 994 | 992 | 989 | 998 |  | 400s |
| Complementary & Alternative Med. | 305 | 499 | 890 | 991 | 989 | 1000 | 1000 | 1000 | 1000 | 1000 | 1000 |  | 500s |
| Dentistry | 279 | 347 | 642 | 596 | 709 | 789 | 830 | 829 | 851 | 867 | 912 |  | 600s |
| Evolutionary Biology | 544 | 898 | 944 | 944 | 931 | 947 | 956 | 982 | 980 | 1000 | 1000 |  | 700s |
| Genetics | 1000 | 1000 | 1000 | 1000 | 1000 | 1000 | 1000 | 992 | 1000 | 1000 | 1000 |  | 800s |
| Human Movement & Sports Science | 403 | 456 | 542 | 749 | 824 | 860 | 990 | 946 | 1000 | 997 | 1000 |  | 900s |
| Immunology | 873 | 921 | 909 | 997 | 967 | 999 | 999 | 976 | 984 | 999 | 1000 |  | |
| Medical Biochemistry & Metabolomics | 331 | 437 | 441 | 599 | 902 | 733 | 714 | 784 | 870 | 999 | 1000 |  | |
| Medical Microbiology | 877 | 907 | 904 | 916 | 935 | 934 | 993 | 972 | 1000 | 999 | 1000 |  | |
| Medical Physiology | 347 | 393 | 647 | 843 | 975 | 996 | 1000 | 1000 | 1000 | 1000 | 1000 |  | |
| Microbiology | 1000 | 1000 | 1000 | 970 | 967 | 1000 | 1000 | 1000 | 1000 | 998 | 1000 |  | |
| Multidisciplinary Journals | 1000 | 1000 | 1000 | 1000 | 1000 | 1000 | 1000 | 1000 | 1000 | 999 | 1000 |  | |
| Neurosciences | 959 | 940 | 933 | 959 | 993 | 994 | 998 | 997 | 998 | 999 | 997 |  | |
| Nursing | 768 | 857 | 941 | 999 | 1000 | 997 | 997 | 996 | 999 | 993 | 1000 |  | |
| Nutrition & Dietetics | 388 | 561 | 711 | 925 | 941 | 964 | 1000 | 1000 | 1000 | 1000 | 1000 |  | |
| Oncology & Carcinogenesis | 996 | 992 | 998 | 998 | 999 | 997 | 1000 | 1000 | 1000 | 998 | 999 |  | |
| Ophthalmology & Optometry | 608 | 698 | 685 | 689 | 815 | 807 | 932 | 899 | 886 | 931 | 988 |  | |
| Paediatrics & Reproductive Med. | 652 | 667 | 818 | 826 | 892 | 914 | 978 | 980 | 987 | 981 | 989 |  | |
| Pharmacology & Pharmaceutical Sci. | 739 | 823 | 836 | 918 | 977 | 959 | 971 | 982 | 997 | 974 | 980 |  | |
| Physiology | 226 | 297 | 434 | 556 | 712 | 799 | 945 | 996 | 999 | 999 | 1000 |  | |
| Unclassified Journals | 851 | 855 | 919 | 958 | 954 | 937 | 943 | 908 | 871 | 893 | 998 |  | |

Values represent the number of papers identified and screened in each field for each year. Light green shades correspond to lower numbers (typically due to a limited number of publications being accessible in PubMed Central for the specified field and year), whereas dark blue shades correspond to higher numbers. Abbreviations: Med., medicine; Sci., science.
